# Evaluating the contribution of aestivation to the persistence of malaria mosquitoes through the Sahelian dry season using stable isotopes

**DOI:** 10.1101/2021.11.17.468867

**Authors:** Roy Faiman, Alpha S. Yaro, Adama Dao, Zana L. Sanogo, Moussa Diallo, Djibril Samake, Ousmane Yossi, Laura R. Veru, Leland C. Graber, Abigail R. Conte, Cedric Kouam, Benjamin J. Krajacich, Tovi Lehmann

## Abstract

Data suggests the malaria vector *Anopheles coluzzii* persists in the Sahel by dry-season aestivation though evidence is scant. We have marked *Anopheles* mosquitoes using deuterium (^2^H) to assess the contribution of aestivation to persistence of mosquitoes through the seven-month dry season. If local aestivation is the only way *A. coluzzii* persists, the frequency of marked mosquitoes should remain stable throughout, whereas finding no marked mosquitoes would be evidence against aestivation. Larval sites were spiked with ^2^H at the end of the 2017 wet season in two Sahelian villages in Mali. We monitored ^2^H-enriched populations until the onset of rains. By the end of the enrichment period, 33% of *A. coluzzii* mosquitoes were clearly marked. Expectedly, ^2^H levels in marked mosquitoes degraded over time, resulting in a partial overlap of the marked and non-marked ^2^H distributions. We utilized three methods to estimate the fraction of marked mosquitoes in the population. Seven months after enrichment, 7% of the population had ^2^H values above the highest pre-enrichment value. An excess of 21% exceeded the 3^rd^ quartile of the pre-enrichment population. A finite mixed population model showed 2.5% represented a subpopulation of marked mosquitoes with elevated ^2^H, compatible with our predictions. We provide evidence that aestivation is a major persistence mechanism of *A. coluzzii* in the Sahel, contributing at least 20% of the adults at the onset of rains, suggesting *A. coluzzii* utilizes multiple persistence strategies enabling its populations rapid buildup, facilitating subsequent malaria resurgence. These may complicate vector control and malaria elimination campaigns.

**Significance statement:** Here we estimated the contribution of aestivation to the persistence of mosquitoes through the seven-month long dry season, by marking a known fraction of the adult population through larval site ^2^H-spiking by the end of the wet season and assessing the change in this fraction through the dry season, until after the onset of the first rain of the subsequent wet season. In a mark-release-recapture study using stable isotopes, we provide compelling evidence that the primary Sahelian malaria vector *Anopheles coluzzii* aestivates on a population-scale, contributing at least 20% of the adults which reestablish the population of the subsequent wet season. The capacity to use multiple strategies of persistence in time and space might complicate vector control and elimination campaigns.

## Introduction

According to the latest World Malaria Report, there were 229 million cases of malaria in 2019 and the estimated number of malaria deaths stood at 409,000. Africa continues to carry a disproportionately high share of the global malaria burden, amounting to 94% of all malaria cases and deaths (World Health Organization, 2020).

In the West-African Sahel, malaria transmission starts following the monsoon rains and cases peak between September and November, when surface water dries out. In the dry season (December-May) rain is scant, surface water is absent, and therefore mosquito densities and disease transmission drop markedly (1, 2). During the dry season *Anopheles coluzzii* has been shown to persist at low frequency, while *A. arabiensis* and *A. gambiae* sensu stricto (hereafter *A. gambiae*) are virtually absent (Lehmann et al. 2010, Adamou et al., 2011; Dao et al., 2014). For a 1-3 weeks period every year, in late March to early April, *A. coluzzii* densities rebound inexplicably (3). The source of the mosquitoes during this brief period has been debated and contended over the years between supporting local aestivation or long-distance migration. During this abrupt 10-20 day period of the dry season, mosquito density spikes to levels resembling or even superseding the wet season maxima, only to vanish again until the first rain that marks the onset of the new rainy season in May-June. Days after the first rain, and importantly *before* a new generation of mosquitoes can develop to adulthood, populations of *A. coluzzii* appear (Lehmann et al 2010, Adamou et al., 2011; Dao et al., 2014). On the other hand, the populations of *A. arabiensis* and *A. gambiae* start the population growth a 6-8 weeks period after the onset of the rains (Adamou et al., 2011; Dao et al., 2014).

To date, direct evidence, such as the capture of aestivating adult mosquitoes has remained limited (4, 5). Although these studies have demonstrated that aestivation does occur, they do not rule out additional strategies such as long-distance migration and have not assessed the relative contribution of aestivation to the persistence of mosquitoes over the dry season.

Tracking small insects over extended time (or large distances) remains a challenge given their small size and methodological limitations in marking large numbers for extended periods (6, 7). Various adult insects retain the deuterium concentration from their larval site in their hardened chitinous tissues (e.g., wings); which was used to determine the provenance of several lepidopteran and Odonata species which migrate over large distances, e.g., the Monarch butterfly, the Globe Slimmer Dragonfly and the painted lady butterfly and (8–10). Mosquitoes developing in [^2^H]-enriched water are marked with an elevated proportion of deuterium (11), providing a new and promising opportunity to test whether *A. coluzzii* persists in the Sahel through the dry season via aestivation. This marking approach potentially enables long-term, mass-marking, albeit with an expected signal degradation or “decay” over a prolonged period as was evident in mosquitoes (Faiman et al., 2019). Hydrogen loss from chitinous tissues was shown to occur in insects and crustaceans in regions of α-chitin, caused by deuterium exchange with protium (^1^H), enzymatic digestion and hydrolysis (12, 13). Here, we aimed to estimate the contribution of aestivation to the persistence of mosquitoes through the 7-month dry season by marking a known fraction of the adult population through natural larval site ^2^H-spiking by the end of the wet season (October-November) and assessing the change in this fraction through the dry season, the late dry season peak (March-April), and especially at the onset of the first rain of the subsequent wet season, before a new generation of mosquitoes may develop. If aestivation is the only way *A. coluzzii* persists, the fraction of marked mosquitoes should remain stable. Finding no marked mosquitoes with adequate sampling or a much smaller fraction of marked mosquitoes would be evidence against aestivation or for low contribution of aestivation, respectively, raising the possibility of alternative strategies.

## RESULTS

### Enrichment during late wet season (September-November)

Variation between villages and species Isotopic enrichment of natural larval sites began in late September when all three anopheline vector species are found in varying proportions. Enrichment ended when all surface waters dried up in late November. To determine the magnitude of adult enrichment and assess the proportion of marked *A. coluzzii* at the time, a comparison of the species composition and their respective enrichment level was done in both focal villages. Species composition in Thierola and M’Piabougou before and during enrichment (September-November) was similar (September: exact test P>0.066, October: exact test P>0.46, November: exact test P>0.99, and overall using Cochran-Mantel-Haenszel Statistics: Nonzero correlation_df=1_= 0.7023, P>0.402, General association_df=2_=3.9075, P>0.95) and therefore villages were pooled (Fig. 1a). The highest ^2^H value of the pre-enrichment distribution was 142.2 ppm (n=66, Fig. 1c). Mosquitoes from enriched larval sites were identified based on ^2^H values above 145 ppm (Fig. 1c and Fig. S3). In Thierola, highly enriched mosquitoes representing marked individuals (^2^H>160 ppm; Fig. 1c) surpassed 40% in October, before any were observed in M’Piabougou (Fig. 1b). By November, 90% of mosquitoes collected indoors in Thierola were highly enriched mosquitoes, but this fraction was near 15% in M’Piabougou (Fig. 1b). Between-species differences in the proportions of marked mosquitoes were detected in both villages with *A. coluzzii* showing lower marking than *A. gambiae* and *A. arabiensis* (Fig. 1b). The effect of experimental enrichment on ^2^H values of mosquitoes collected indoors in each village is evident by the large gap between natural ^2^H values and the values of some of the mosquitoes after enrichment began (Fig. 1c). The increase in ^2^H enrichment levels over time in mosquitoes (November vs. October) reflects the increasing concentration of ^2^H in the larval sites (see Methods).

**Figure 1.**
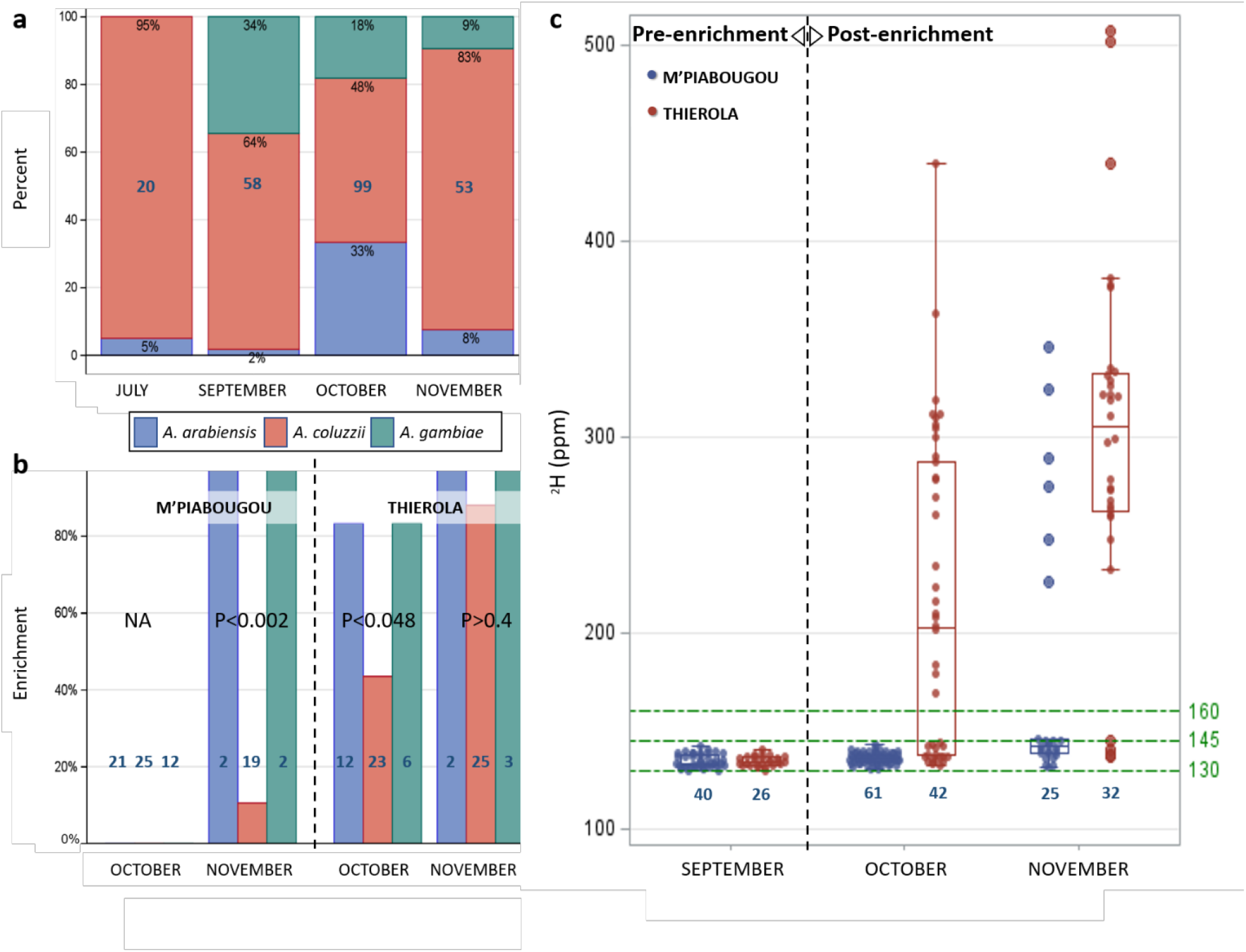
**a**. Species composition of indoor collections in Thierola and M’Piabougou (pooled) before (July and September) and during the enrichment (October-November). Sample size (blue) and proportion of each species (black) are shown inside bars (percent). Samples were taken from both villages with minimum sample size/village of n=23, except in July (Thierola: n=20). Differences in composition between villages were not significant: September (exact test P>0.071), October (exact test P>0.46), November (exact test P>0.99) and overall using Cochran-Mantel-Haenszel Statistics (Nonzero correlation/Row mean score difference_df=1_= 0.7023 P>0.402, General association_df=2_=3.9075, P>0.95). **b**. The change in the proportion of marked mosquitoes (^2^H>160 ppm) after enrichment (October-November) in each village by species (colors). Exact tests of heterogeneity among species were run using Monte Carlo estimation (10,000 pseudo samples). Sample size is given in blue. **c**. Distribution of ^2^H values in each village before (September) and during enrichment (October-November) by village. Sample sizes are given in blue and reference lines (green, dot-dashed) delimit natural variation and the gap between them and experimentally enriched (marked) mosquitoes (145-160 ppm).

The exceptionally high proportion of marked adults in Thierola (Fig. 1b) is attributed to the fact that during the late rainy season, the village has only two major larval sites; one bordering the village from the west and the other, located 1.2 km NE of the nearest house, as opposed to more than 15 larval sites in M’Piabougou (Fig. S2), where enrichment started later and progressed slower than in Thierola. Additionally, most mosquitoes in Thierola collected indoors were from houses near the main larval site, whereas the collections in M’Piabougou were scattered across the village. In Thierola, the proportion of marked mosquitoes was 92% in the houses flanking the larval site, ∼10-50 m from the water’s edge (n=36), 60% in houses ∼50-150 m away (n=30), and 20% in the houses ∼150-600 m away (n=5) (χ^2^_df:2_=16.5, P<0.001), reflecting the accumulation of newly emerged marked mosquitoes in the houses nearest to the larval site and their slower dispersion across the village. Accordingly, the adjusted estimate of the fraction of marked *A. coluzzii* mosquitoes in Thierola in November, based on the number of houses per area was 56.4%. In M’Piabougou, on the other hand, the comparable proportion of marked *A. coluzzii* reached 10%.

### Tracking marked mosquitoes during the dry season (December-April)

The first half of the dry season (December-February) is characterized by few mosquitoes found indoors, all of which are *A. coluzzii* (3). To avoid impinging on the aestivating population only a limited number of anophelines were collected to assess enrichment levels. Because of the expected attrition in ^2^H over 4-6 months since marking (11) and (12, 13), we hypothesized that certain marked mosquitoes which their ^2^H values degraded enough may “accumulate” in the upper tail of the population’s ^2^H distribution, thus inflating the expected percentile. Accordingly, the proportion of mosquitoes in the post-enrichment population whose ^2^H values were above the 3^rd^ quartile of the pre-enrichment distribution would be larger than the expected 25% (Methods). After the last larval site dried up in Thierola and in M’Piabougou (see Material and Methods), no rain fell in the area during the dry season and no surface water was available for larvae. Therefore, any mosquito with a ^2^H value above natural levels is considered an aestivator marked during the previous wet season. Due to the low numbers of mosquitoes in the villages during December-February, only 7 were collected (6 from Thierola); all were *A. coluzzii*. As the late-dry-season peak unfolded in March-April (14), an additional 102 specimens were collected from Thierola, all *A. coluzzii* (n=100, 2 failed to produce a PCR band), in accord with previous results on seasonal species composition (3, 4, 14, 23).

During the early dry season (December to February), 1-3 months after marking, the ^2^H values of mosquitoes were lower than those during the enrichment period (October-November), but two of the seven mosquitoes exceeded the highest ^2^H values measured in the pre-enrichment levels (142 ppm, Fig. 2a), suggesting that they were enriched or marked mosquitoes. Therefore, approximately, one third of the mosquitoes in December-February from Thierola (Fig. 2a, ^2^H >152 ppm, one from mid-December and another from the end of January) were, therefore, considered as aestivators, or 29% from the focal villages (2 of 7 mosquitoes).

**Figure 2.**
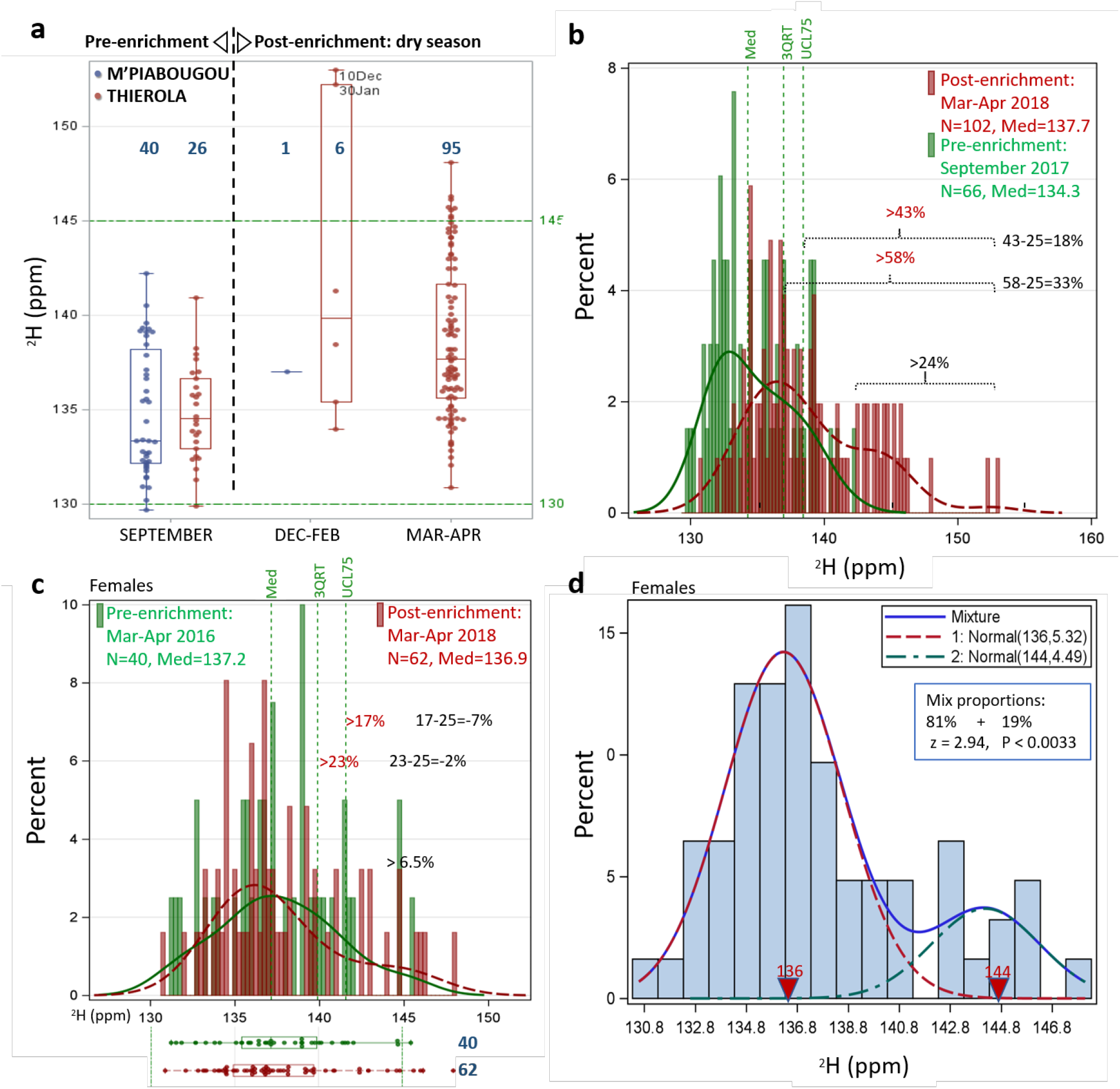
Evaluation of the proportion of marked mosquitoes during the dry season by different methods. **a**. Comparing the post-enrichment distribution of December-February and March-April 2018 with pre-enrichment (September 2017) in Thierola and M’Piabougou by box-whisker plots; **b**. Comparing the post-enrichment distribution of March-April 2018 with pre-enrichment (September 2017) in Thierola by histogram showing the median (Med) and 3^rd^ quartile (3QRT) as well as the 95% upper confidence limit (UCL75) of the 3^rd^ quartile (green dashed vertical lines) of the pre-enrichment distribution. The expected proportion of the distribution beyond the maximum, 3^rd^ quartile and upper 95% upper confidence limit of the 3^rd^ quartile are shown above the horizontal braces (see text); **c**. Comparing the post-enrichment distribution of March-April 2018 with March-April 2016 in Thierola (females only) using box whiskers (bottom) and histogram (top) and employing the quartile method; **d**. The composition of the post-enrichment females (n=62) using finite mixed distribution model assuming two subpopulations (initial values of mean and variance 134 ppm and 8 ppm^2^, and 144 ppm and 5 ppm^2^, for each subpopulation, respectively (see Methods). The results support a mixed population with of a mean of 136.3 ppm and variance of 5.7 ppm^2^ (P<0.001, 81%), and a smaller subpopulation of a mean of 144.2 ppm and variance of 4.5 ppm^2^ (P<0.001, (19%), which are significantly separated (P<0.0033).

During March-April, 24% of the mosquitoes collected in Thierola exceeded the highest ^2^H value of the pre-enrichment period (142 ppm, Figs. 2a and 2b) and ∼10% surpassed 145 ppm. Because of the expected “decay” in ^2^H over 4-6 months since marking we predicted there should be an excess of mosquitoes in the upper end of the distribution (see Methods). As predicted, the proportion of mosquitoes above the 3^rd^ quartile of the pre-enrichment distribution was 58%, indicating an excess of 33%. Moreover, the proportion of mosquitoes in the post-enrichment distribution that had values above the 95% upper confidence limit (UCL) of the 3^rd^ quartile (of the pre-enrichment distribution) was 43%, indicating a significant excess of at least 18% (Fig. 2b).

The shrinking gap between the pre- and post-enrichment values raised the possibility that seasonal variation in natural ^2^H might confound the results. Therefore, we compared the post-enrichment distribution in March-April 2018 with the same period in 2016 (Fig. 2c). Because only females were available from the 2016 collections, we excluded males from the 2018 sample. Notably, the ^2^H distribution of March-April 2016 had higher median and maximum values (137.2 ppm, 146 ppm, Figs. 2b, 2c) than the corresponding values of the pre-enrichment distribution of September 2017 (134.3 ppm 142.2 ppm) and even the post enrichment distribution of March-April 2018 (136.9 ppm, Fig. 2c), suggesting a seasonal and/or inter-annual variation in natural ^2^H. Likewise, a lower proportion of post-enrichment mosquitoes exceeded the maximum of the 2016 distribution (6.5%, Fig. 2c) and no evidence was found for an excess of post-enrichment mosquitoes above the 2016’s 3^rd^ quartile (−2%, Fig. 2c).

We used the finite mixed distribution model (FMM; see Methods) as a reference-free approach to evaluate the presence and composition of the March-April population. The result for females (n=62) indicated the mixture of two subpopulations, with the larger subpopulation (81%) with a mean of 136.3 ppm (P<0.001) and variance of 5.7 ppm^2^, and a smaller subpopulation (19%) with a mean of 144.2 ppm (P<0.001) and variance of 4.5 ppm^2^, which were statistically separate (P<0.0033, Fig. 2d).

### Tracking marked mosquitoes during the onset of rain (May-June 2018)

The first rain falls typically between mid-May to mid-June, and the first larvae are spotted in the newly formed larval sites 10-20 days after the first rain (4, 14). The first rain of 2018 fell on June 24^th^. Hence, mosquitoes collected from May 15 to July 5^th^ represent adults that in high probability were not mixed with the young adults which developed in the newly formed larval sites. Although few putative aestivators may remain for a couple of additional weeks, their proportion is expected to decrease rapidly. Accordingly, we assumed that mosquitoes collected from July 17^th^ to July 30^th^ represented mostly the new cohort of adults which developed in the freshly formed larval sites. All mosquitoes tested from M’Piabougou were *A. coluzzii* (100%, n=107), as were 99% (n=186) from Thierola, in agreement with previous results (3, 4, 14, 23). The first *A. arabiensis* and *A. gambiae* were detected in Thierola on June 22^nd^ and July 5^th^, respectively.

Although lower than during the late dry-season peak (March-April), the ^2^H values of mosquitoes from Thierola and M’Piabougou at the onset of rains (May 15^th^-July 5^th^) were higher than the pre-enrichment period (September 2017) and their right tail exceeded the highest ^2^H values measured in the pre-enrichment population (Fig. 3a). As expected, based on aestivation, three weeks later, the ^2^H median and 3^rd^ quartile dropped in both villages (median difference of 1.6 ppm, P<0.0023 t_1_=3.1; 3^rd^ quartile difference of 2.2 ppm P<0.0001, t_1_=4.0; 0.9 quantile difference of 3.4 ppm P<0.022, t_1_=2.3, quantile regression), representing the first generation of mosquitoes which developed in the newly formed larval sites (late July). Notably, the difference increased at higher quantiles (P<0.052, χ^2^_2_=5.94) indicating that the greatest difference was related to highest ^2^H values. As expected, the ^2^H range of this first cohort of the 2018 wet season was similar to that of the pre-enrichment period (September 2017, median difference of 0.5 ppm, P>0.53 t_1_=0.7; 3^rd^ quartile difference of 0.4 ppm P>0.4, t_1_=0.5; 0.9 quantile difference of -1.5 ppm P>0.09, t_1_=-1.7, quantile regression). The difference between Thierola and M’Piabougou was minimal and non-significant (median difference of 0.42 ppm, P>0.31 t_1_=1.0; 3^rd^ quartile difference of 0.6 ppm P>0.5, t_1_=1.2; 0.9 quantile difference of 0.75 ppm P>0.4, t_1_=0.8, quantile regression), thus, we pooled mosquitoes from these villages at this time point. In May-July, 6.5% of the population have exceeded the highest value in September (142.2 ppm, Figs. 3a and 3b; four mosquitoes exceeded the 145 ppm mark).

**Figure 3.**
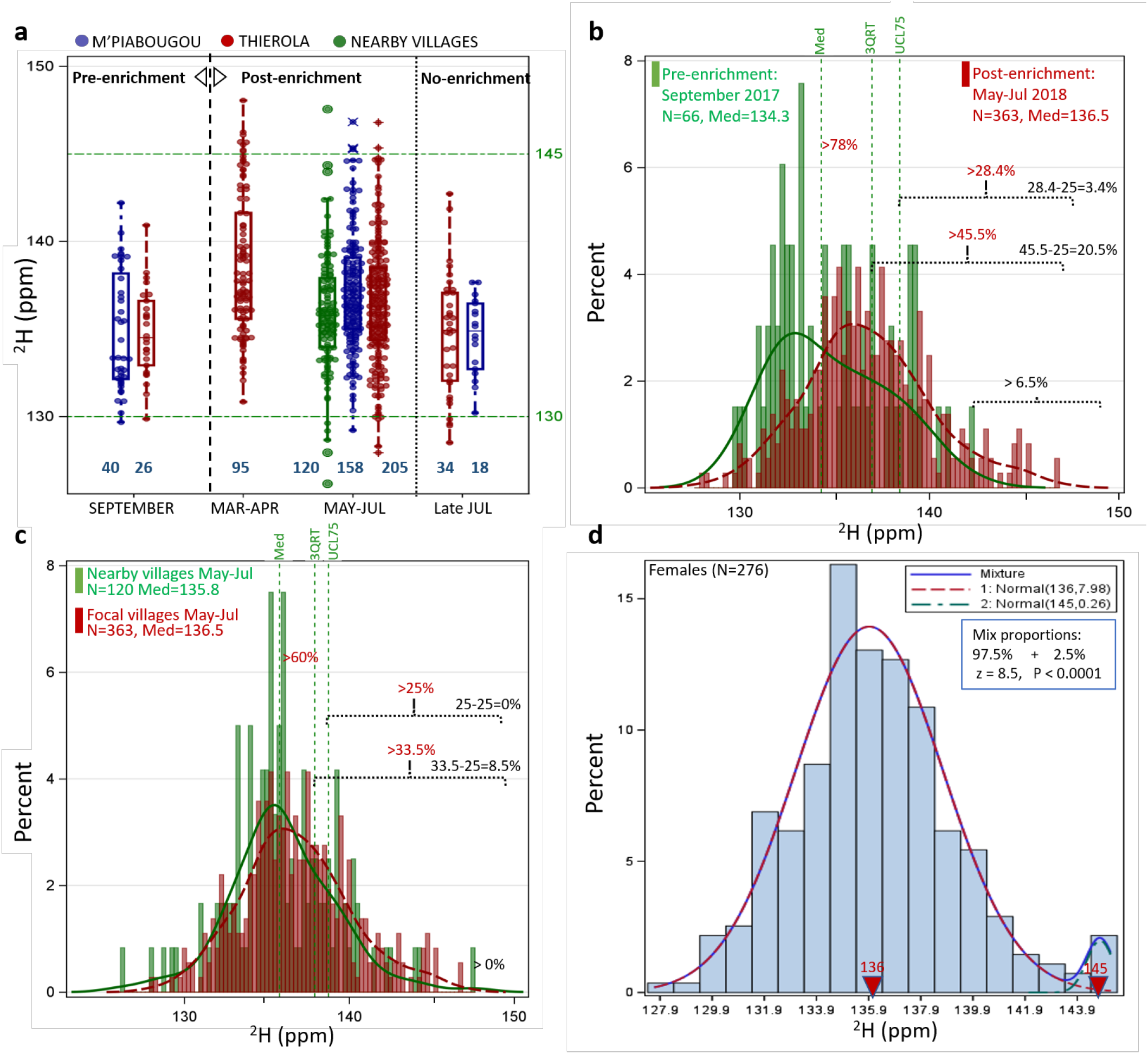
Evaluation of the proportion of marked mosquitoes during the onset of rains, 7-8 months after end of enrichment. **a**. Comparison of the pre-enrichment (September 2017) in Thierola (red), M’Piabougou (blue) and nearby villages (3-7 km) (green) with corresponding post-enrichment distribution of March-April 2018 and May-Jul 2018 (May 15-July 5), as well as the late July 2018 (July 18-30, reflecting first cohort of mosquitoes growing in newly formed larval sites the following season – ‘No enrichment’) by box-whisker plots; **b**. Comparing the post-enrichment distribution of May-July 2018 with pre-enrichment (September 2017) by histogram showing the median (p50) and 3^rd^ quartile (p75) as explained in Fig. 2; **c**. Comparing the post-enrichment distributions of May-July 2018 in focal villages and nearby villages (3-7 km away) and employing the quartile method described above; **d**. The composition of the May-July post-enrichment females (n=276) using finite mixed distribution model (FMM with two subpopulations (initial mean and variance 134 and 8, and 144 and 5, for each subpopulation, respectively). The results support a mixed population with a mean of 136.3 ppm and variance of 5.7 ppm^2^ (P<0.001, 81%), and a smaller subpopulation of a mean of 144.2 ppm and variance of 4.5 ppm^2^ (P<0.001, 19%), which are significantly separated (P<0.0033).

Assessment of excess mosquitoes whose ^2^H values were above the 3^rd^ quartile of the pre-enrichment population (see Methods and above) revealed 45.5% of post-enrichment mosquitoes, indicating an excess of 21%. Moreover, 3.4% of the post-enrichment mosquitoes represented excess over the 95% UCL of the 3^rd^ quartile of the pre-enrichment distribution, i.e. a statistically significant excess (see Methods). To further evaluate the presence of enriched mosquitoes in the experimental (or focal) villages with respect to seasonal and or inter-annual change, we compared the post-enrichment population with the synchronous population (May-July 2018) in four villages 3-7 km from the experimental villages (Zanga, Bako, Dodougou, and Dougouguele, Figs. S2, 3a, 3c). The median of the post enrichment May-July 2018 was higher than that of the nearby villages (Figs. 3a, 3c, P<0.02 quantile regression). Although the maximum value of the ^2^H distribution of the population during the onset of rains was lower than that of the nearby villages (Fig. 3c), the proportion of mosquitoes that exceeded the third quartile of the ^2^H distribution of the nearby villages was 33.5%, yielding an excess of 8.5%. Notably, there were no mosquitoes that exceeded the 95% UCL of the 3^rd^ quartile (0%, Fig. 3c).

Using FMM to evaluate the homogeneity of the female populations during the onset of rains, we detected evidence for a mixed population with the larger subpopulation (97.5%) with a mean of 136 ppm (P<0.001) and variance of 8 ppm^2^, and a smaller subpopulation (2.5%) with a mean of 145 ppm (P<0.001) and variance of 0.3 ppm^2^, which were significantly separated (P<0.0001, Fig. 3d).

### Alternative sources of mosquitoes with elevated ^2^H values in Thierola

The shrinking gap between pre- and post-enrichment mosquitoes 4-8 months after end of marking and the evidence for seasonal and/or inter-annual variation in ^2^H (Fig. 2c) highlighted the need to characterize the spatio-temporal variability in natural ^2^H. We used mosquito samples representing natural ^2^H across seasons and years from Thierola, nearby (3-7 km) villages, and distant villages (25-120 km) away (Fig. S1). Using quantile regression (Methods, Supp. Table S1), we estimated seasonal variation at the different sites across quartiles (Fig. 4a). The results revealed a geographical cline in mosquitoes with increasing ^2^H values towards the north and a seasonal fluctuation in ^2^H levels in each location, which was more pronounced in the typical Sahelian villages (4 ppm range) compared with the rice irrigation area of Niono (2.5 ppm range, Fig. 4a). Although seasonal dynamics of ^2^H levels showed site-specific pattern, a “convergence” across sites was observed in March-April – the peak ^2^H level in Thierola (and Ballabougou, Fig. 4a). To further assess if Niono could serve as the source of the migrants to Thierola (and Ballabougou), confounding the post-enrichment distributions in March-April and/or May-June, these distributions were compared (Figs. 4b and 4c). In March-April these populations exhibited similar distributions, except mild excess of higher ^2^H values in Thierola (Fig. 4c). However, in May-June the differences were pronounced, precluding the possibility of mass migration from Niono into Thierola.

**Figure 4.**
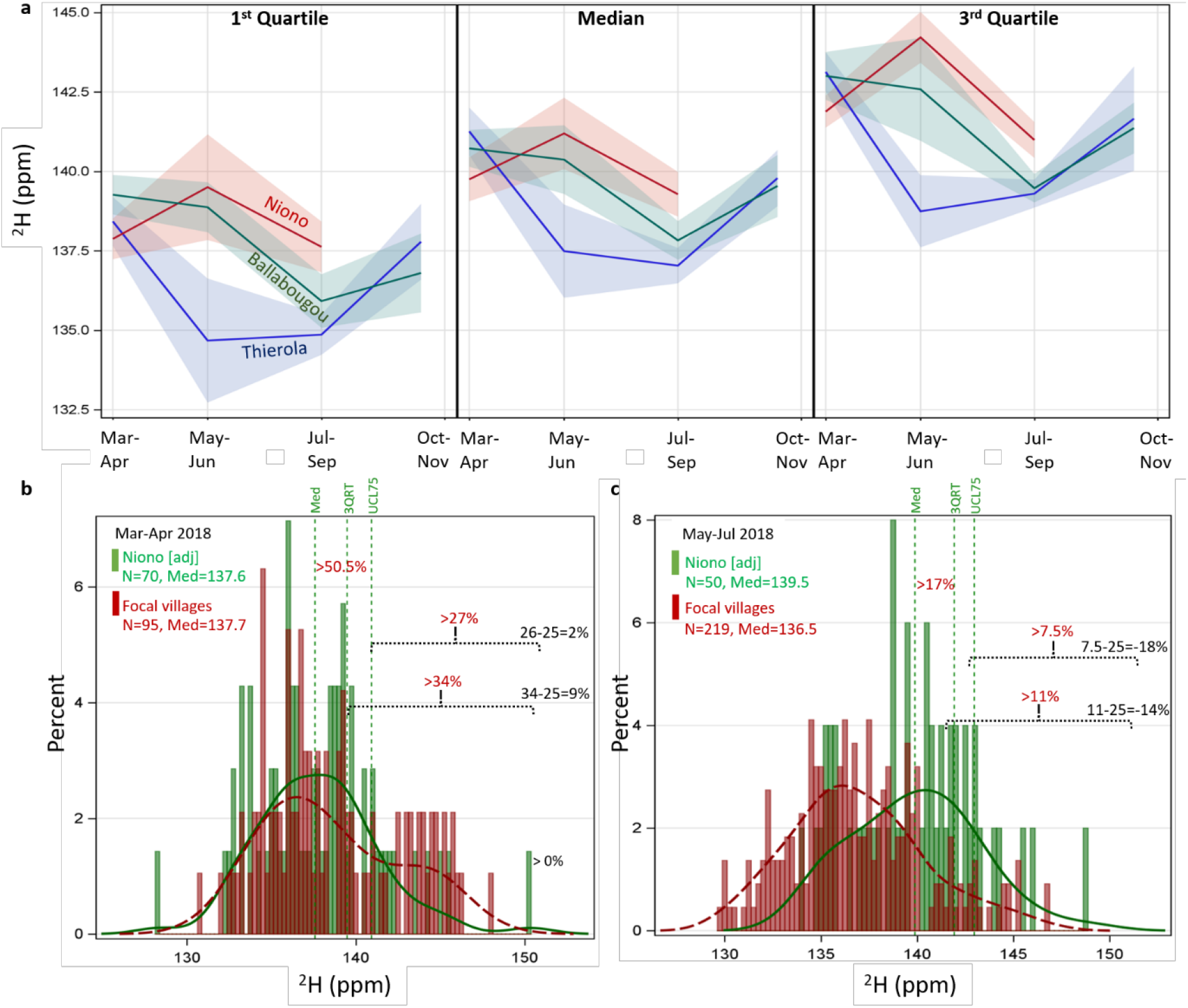
Spatial and seasonal natural variation in ^2^H and evaluating the possibility that mass migration from Niono accounts for the elevated ^2^H values in post-enrichment Thierola. **a**. Seasonal natural variation in ^2^H in Thierola, Ballabougou, and Niono evaluate over the quartiles of the distributions using quantile regression (Table S1). Bands represent 90% confidence intervals. Note: during October-November, no sample was available from Niono. Evaluating the similarity between the post-enrichment populations from Thierola and the concomitant population from Niono using histograms showing the median (p50) and 3^rd^ quartile (p75, as explained in Fig. 2) in March-April **(b)** and in May-June **(c)**.

## Discussion

This novel mosquito mark-capture study, using stable isotopes, was aimed to evaluate the contribution of aestivation to the persistence of Sahelian *Anopheles* throughout the dry season by the ultimate evidence – tracking mosquitoes marked at the end of the wet season – until the beginning of the following wet season. By the end of the enrichment phase (November 2017), the proportion of *A. coluzzii* marked in Thierola and M’Piabougou was 56% and 10% respectively, and even higher percentages of *A. gambiae* and *A. arabiensis*. This difference between species is consistent with previous results suggesting that *A. coluzzii* density plummets indoors and in larval sites during this period, presumably as *A. coluzzii* depart to some undiscovered shelters and/or prepare for aestivation by sugar feeding (14, 24, 25), while *A. gambiae* and *A. arabiensis* attain their indoor peak densities (3, 4, 14). Several months post enrichment, the ^2^H levels in the focal villages had declined as previously described (11) and although some values above pre-enrichment levels persisted until July 2018, estimating the proportion of ^2^H marked mosquitoes required use of multiple methods. Early after the end of enrichment (December-January), a small sample available from Thierola (n=7) revealed that 29% of the *A. coluzzii* population in the combined focal villages were enriched. At the onset of rains of the next wet season (May-June 2018), seven months after enrichment, 7% of the population had ^2^H values above the pre-enrichment maximum and the excess over the 3^rd^ quartile of the pre-enrichment population was 21% (September 2017, Figs. 3a and 3b). Concomitant comparison with the neighboring villages (3-7 km away) revealed 9% and 10% excess over the 75^th^ and the 50^th^ percentiles (i.e., 3^rd^ quartile and median), respectively, although only the difference between the medians was significant (P<0.02 quantile regression, Figs. 4a and 4c). Finally, FMM analysis supported a mixed population with 2.5% representing a subpopulation with elevated mean ^2^H compatible with our predictions (Fig. 4d). The consistent trend of decline in the estimates of aestivators add credence to the results and hints to the role of dispersal of aestivators among neighboring villages (below). Altogether, these results establish the presence of mosquitoes with elevated ^2^H levels over that expected naturally, i.e., marked mosquitoes, and thus support local persistence of a significant part if not all *A. coluzzii* over the long dry season in the Sahel.

Notably, the results for the late dry-season peak (March-April 2018) provided weaker support for aestivation, despite finding that 24% exceeded the maximum of the pre-enrichment period (10% >145 ppm, Figs. 2a and 2b) and 33% represented excess over the 3^rd^ quartile of the pre-enrichment distribution (Fig. 2b), because comparison with March-April 2016 suggested that a seasonal variation in ^2^H levels could also have given rise to a similar distribution. In this comparison, 6.5% of post-enrichment mosquitoes exceeded the maximum of the 2016 distribution but none exceeded the 2016 3^rd^ quartile (Fig. 2c). The results of the FMM provided further support for aestivation by indicating a mixed (female) population with 19% with a mean of 144.2 ppm (P<0.001) which were statistically separated P<0.0033 (Fig. 2d), yet migrants at this period may also generate a similar outcome (below). Therefore, these results cannot rule out alternative hypotheses, such as migration, in explaining the ^2^H distribution of March-April 2018.

The decay of ^2^H values in field- ^2^H-enriched mosquitoes over 4-8 months (11) and the realization of a spatio-temporal variation in natural ^2^H level led us to use multiple approaches to evaluate the proportion of marked mosquitoes with elevated ^2^H values in the focal villages. Taking the mean value of these estimates (and pooling the focal villages) for each period to assess the temporal trend revealed that during the late dry-season peak (March-April), 17% of the *A. coluzzii* were marked (assuming no long-range migration), and 6.1% were still marked by the onset of the rains of the subsequent wet season (May 15 – July 5, table S1). Given a starting fraction of 33% (above and Table S1), these results entail that 18% of the mosquitoes at the onset of the new rains were aestivators that survived the 7-8 months long dry season (Fig. 5). This trend indicates an accelerated decline in marked mosquitoes after February (Fig. 5), which may be accounted for by increased dilution due to migration (below). Previous evidence based on mosquito hotspots in Thierola during the dry and wet seasons suggested that mosquitoes shelter outside the boundary of the village and probably within one or a few kilometers from the village (25). Thus, movement between neighboring villages would result in underestimating the actual fraction of aestivators because of the influx of unmarked-immigrant aestivators from neighboring villages and the loss of emigrant-marked mosquitoes who moved into neighboring villages. Indeed, occasional movement between neighboring villages was documented in previous studies in our area (4, 26) and elsewhere in the Sahel (27, 28). The mean and median ^2^H values of the four surrounding villages (3-7 km from the focal villages, Fig. S1) were significantly lower than those of the focal villages (Figs. 3a and 3c), yet the mosquito with the highest ^2^H value at that period was found in Zanga, 3 km north of Thierola (Fig. 3a), consistent with migration among these villages. The accelerated reduction in marked mosquitoes (Fig. 5) after January coincides with the period of change in the village hot-spots that supported local movement of mosquitoes during the late dry-season in Thierola (25). Likewise, FMM analysis on the females in nearby villages (Zanga, Bako, Dougouguele and Dodougou) supported two populations, with the smaller population consisting of 1% (not shown), as expected if some migrants from the focal villages dispersed to the surrounding villages. Accordingly, the estimate of 18% represents the minimum proportion of aestivators and whether the actual estimate. reaches 100% or an intermediate proportion remains to be elucidated.

**Figure 5.**
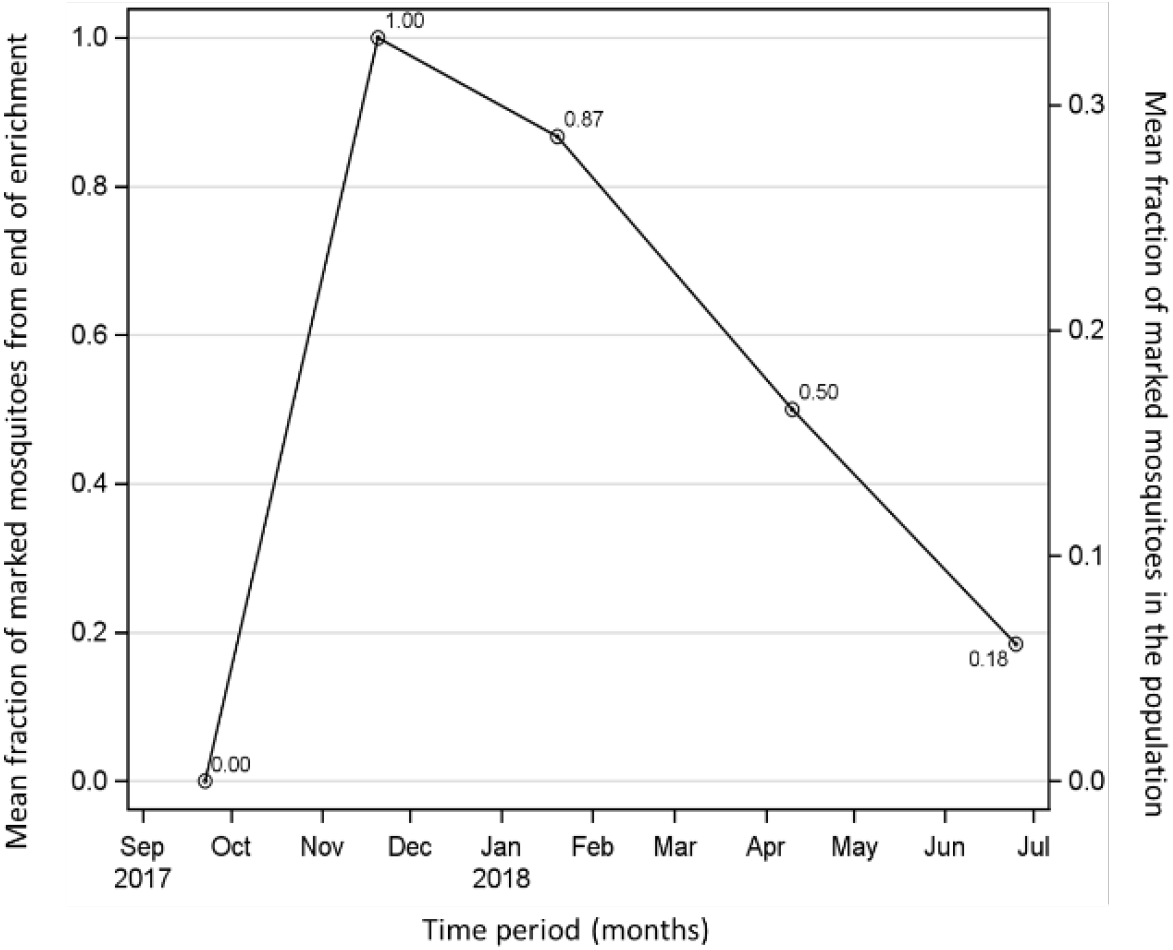
Change in the proportion of marked *A. coluzzii* over the course of the experiment. Values represent the mean fraction over all estimates (Table S1). Right axis expresses proportions of marked mosquitoes from the total sample and the left axis expresses the corresponding values from the end of the marking period for (November 2017, i.e., 0.33 is the 100%). Fraction values of marked mosquitoes from the end of enrichment are given near each point.

Considering the hypothesis that the elevated ^2^H values can be explained without marked-aestivating mosquitoes either by (1) seasonal change in ^2^H in focal (and other Sahelian) populations, and (2) mass-migration from distant localities where background ^2^H levels are higher (9, 29, 30) we have included additional comparisons. In the absence of monsoon rains during the dry season, evoporative-transpiration of the common hydrogen occurs at a higher rate than that of deuterium, resulting in seasonal fluctuation of natural ^2^H values (31–35). Variation in natural ^2^H in Thierola, Ballabougou and Sokourani (Niono) using the median ^2^H followed accordingly (Fig. S4). Although a plausible hypothesis, the lower median ^2^H in neighboring villages compared with the focal villages (Fig. 3a and 3c) does not reconcile with either hypothesis because the neighboring villages would be expected to exhibit similar ^2^H levels if either hypothesis was true. The migration hypothesis also requires mosquitoes to fly tens or hundreds of kilometers downwind from an area of exceptionally high density (3, 14). To the best of our knowledge, only the rice irrigation region of Niono (∼140 km NE, or the inner Niger delta near Mopti, ∼300 km NE) meets these requisites during March-April (36, 37), but not during May-June, when the dominant wind direction blows from the South (38). Importantly, natural ^2^H distribution in Thierola was similar to that of Niono only during the late dry season peak (March-April, Fig. 4a), lending support for long-range migration at that period. However, no *A. coluzzii* was caught in high altitude despite considerable effort to intercept them, whereas 24 were intercepted in altitude between late July to November, 100-300 m above ground between 2013 and 2015 (39), and additional unpublished data).

To further evaluate the role of long-range migration based on the natural ^2^H distributions, we used samples taken from Thierola over 4 years and examined the seasonal change in the following parameters: i) between-sample median ^2^H, ii) within-sample spread (IQR) and iii), within-sample asymmetry (skewness). Unless the fraction of immigrants is negligible or very large, a mixed population with migrants from distant northern population(s) is expected to exhibit a higher median, larger spread, and positive skewness. With increasing proportion of migrants in the population, the center of the ^2^H distribution (median) will shift upwards, while the effect on the tails of the distribution will be reversed and diminished. If the whole population consists of migrants from a distant northern population, the only change that will be detected is a sharp increase in the median. The samples from Thierola suggest a sharp increase in median ^2^H in March-April, without change in spread and with minimal positive skewness (Fig. S4), compatible with a population consisted of migrants from a northern source (such as Niono). Notably, during July-September the median and the spread were low, but skewness was negative in all four samples from July and in one of five September samples, consistent with an influx of immigrants from the south, during the period of predominant southerly winds and when high altitude migrants were detected (38, 39). Possibly the marked negative skewness in September 2018, may have reflected the exceptional delay in rainfall and the resulting smaller local populations, resulting in a higher proportion of migrants (Figs. 5 and S4).

These mark-capture results including the persistent decline in the fraction of aestivators detected throughout the dry season (Fig. 5) provide compelling evidence that aestivation contributes at least 20% to the persistence of *A. coluzzii* until the onset of rains. Adamou et al. (3) reached a similar conclusion based on their experiment using selective removal of dry-season mosquitoes. On the other hand, our evidence for aestivating mosquitoes being the source of the late dry season peak remains inconclusive because it cannot rule out migration from distant northern areas. Adamou et al. (2011) have interpreted their results as evidence for aestivation as the primary if not the only source of the dry season peak. Recent results (Djibril et al. unpublished) revealed that the mosquitoes during the dry season and the onset of the rains are rarely infected with *Plasmodium* despite their aggressive biting during the late-dry season peak. The low infection despite moderate carriage of gametocytes in the human residents of the villages is more simply explained by migrants than by aestivators because the irrigated areas are known for low mosquito infection rate and because long-distance migrants are expected to consist of young mosquitoes that will not survive long in the Sahel during the dry season. Nevertheless, these explanations are not mutually exclusive, and the population may consist of both aestivators and migrants at that time.

## Conclusions

Our results, based on mark recapture data, provided direct evidence that aestivation is a mechanism that facilitates the persistence of *A. coluzzii* in the Sahel where it contributes at least 20% of the adults that appear near the onset of rains. However, the source of the mosquitoes that appear during the late dry-season peak (March-April) remains unknown because our results support reasonably well both aestivation and long-distance migration from northern areas such as Niono. The capacity to use multiple strategies of persistence in time and space might complicate vector control and elimination campaigns.

## Materials and Methods

### Study area

The field-work including marking and mosquito collection was conducted in the Sahelian villages of Thierola (−7.215 E, 13.658 N) and M’Piabougou (−7.191 E, 13.599 N), Mali, which are 6 km apart and are referred to as the focal villages (Fig. S1). Collections of adult mosquitoes, without marking by ^2^H spiking were also conducted in the neighboring villages (3-7 km from Thierola): Zanga (−7.220 E, 13.686 N), Bako (−7.265 E, 13.643 N), Dodougou (−7.160 E, 13.646 N) and Dougouguele (−7.170 E, 13.611 N, Fig. S1). To evaluate geographical variation in ^2^H background levels (natural) of adult mosquitoes in the region, indoor collections were also conducted in two distant villages: Ballabougou (−7.386 E, 13.860 N), 22 km NNE of Thierola and Sokourani (−6.050 E, 14.217 N, Fig. S1), 140 km ENE of Thierola. Sokourani is located at the Niono rice irrigation area and is referred to as Niono hence forth. To address spatio-temporal variation in natural ^2^H we also used samples of mosquitoes collected on other studies in Thierola and several other villages between 2015-2019.

### Stable isotope enrichment of larval sites

Enrichment of mosquito larval sites in Thierola and M’Piabougou began on September 23^rd^, 2017 and September 30^th^, 2017 respectively. Enrichment ended November 20^th^, 2017 (Thierola) and December 3^rd^, 2017 (M’Piabougou), when larval sites dried up. The enrichment protocol followed (11). Briefly, 27 larval sites in the focal villages were used for enrichment; some were sections of a larger body of water (natural larval site) that were separated by embankment from the rest of the larval site (below). Medium and large natural larval sites were selected such that they would retain rainwater for the longest period possible, and a minimum of three weeks. Sandbag embankments were erected within the larger larval sites to prevent dilution of enrichment material in the total water volume and ensure the target concentrations could be maintained within the compartments. As larval site water levels receded with time, the embankment locations were adjusted and new ones were erected, increasing the relative area of enrichment and leading to natural increases in ^2^H due to evaporation. We used ^2^H-deuterium-oxide (99.6% D_2_O, Cambridge Isotope Laboratories, Inc. Tewksbury, MA) (also: “Heavy water”, ^2^H_2_O or D_2_O; d2H or ^2^H in short) for larval site enrichment. Initial enrichment using 99.6% D_2_O aimed to reach 2.5ml ^2^H/L, based on the volume of the water measured using maximum and minimum diameters of the larval site and multiple stakes (as dipsticks) to measure average depth (11). Additionally, we supplemented 0.25% (0.625 ml ^2^H/L) on a weekly basis to compensate for underground water infiltration. Every three days we also added 200 ml of a microbiota culture produced from microorganisms harvested from the larval sites and cultured in 1% deuterated larval site water, as ^2^H-enriched larval diet (11). Larval site water volume was measured daily and after each rain compensation dosage of ^2^H was added per the volume of water that was added by the rain at the same concentration (2.5ml ^2^H/L).

### Adult mosquito collection, preservation, and identification

Adult mosquitoes were collected indoors, in both Thierola and M’Piabougou in pre-selected periods: September (mid-wet season) – pre-enrichment, October-November 2017 - enrichment phase (late wet season), December-January (early dry season), March-April 2018 - late dry season peak (late dry season), May-June 2018 - onset of rains of the new rainy season, and July – the first cohort of mosquitoes that developed in the newly formed larval sites after the rain. However, because of the late arrival of the first rains in 2018 (June 24), all mosquitoes collected up to the 10-day period following the rain were considered to be either aestivating or migrating adults because there was no time for a new cohort to be produced (3, 4, 14). Thus, all mosquitoes collected from May 15 until July 5^th^ were pooled to represent the emergence of putative aestivators (or long-distance migrants). The first cohort of mosquitoes which developed in the newly formed larval sites after the rains were collected July 18-25 to ensure minimal overlap between this cohort and the previous population.

Adult mosquitoes (*Anopheles* sp.) were collected indoors in the morning, using a mouth aspirator, within ∼100 houses per village. Collected mosquitoes were identified to species or genus, separated by sex and gonotrophic state before being killed and stored individually over a thin layer of cotton placed over desiccant (Silica gel orange, Cat. No. 10087. Sigma-Aldrich, St. Louis, MO) in 0.6 ml microcentrifuge tubes. On a few occasions we used previously collected samples preserved in 75% ethanol.

Following morphological identification (15), species identity of the members of the *Anopheles gambiae* s.l. complex was resolved using a PCR with two legs as template (16). This assay enabled identification of *A. coluzzii* (formerly *A. gambiae* M form), *A. gambiae* s.s. (formerly *A. gambiae* S form), and *A. arabiensis* without the need for restriction enzymes digest of the PCR product.

### Stable isotope analyses

Quantification of ^2^H-enrichment in mosquitoes was previously described in (11). Briefly, desiccated *A. gambiae* s.l. carcasses were divided to main body parts (head, thorax, abdomen, legs and wings). Only thoraces were included in our analysis as they constituted the largest chitinous mass in the mosquito. The other body parts were either used for molecular taxonomy (legs) or kept for future analyses. In a handful of cases when thoraces were unavailable, a combination of head, wing and legs was used instead. Dry mosquito samples were prepared for IRMS as follows: mosquito samples from either the laboratory or the field experiments (and respective standards; see below) were exposed to the IRMS facility’s laboratory environment for at least 72 hrs prior to analysis, to facilitate equilibration of exchangeable hydrogen with local atmospheric water vapor. Water-vapor equilibrated mosquito thoraces were individually weighed in 8×5 mm silver capsules (Cat. No. D2008. EA Consumables, Inc. Pennsauken, NJ, USA) (mean: 0.25 mg, range: 0.15-0.4 mg). Crushed capsules were interspersed with appropriate standards and blanks approximately every ten samples. Stable hydrogen isotope analyses of wild mosquitos were performed at the Smithsonian Museum Conservation Institute Stable Isotope Mass Spectrometry reference laboratory in Suitland, Maryland, USA on a Thermo Delta V Advantage continuous-flow isotope-ratio mass spectrometer coupled to a Thermo Temperature Conversion Elemental Analyzer via a Conflo IV interface.

Samples were thermally decomposed to H_2_ in a ceramic reactor filled with chromium powder at 1100°C. Non-exchangeable δ^2^H values (i.e. δ^2^H_non-ex_) were determined via 3-point linear calibration (Wassenaar & Hobson, 2003) with three keratin reference materials dispersed every 10 samples: CBS (Caribou Hoof Standard, δ^2^H_non-ex_= -197.0 ‰), USGS42 (Tibetan Human Hair, δ^2^H_non-ex_ = -78.5 ‰), and KHS (Kudu Horn Standard, δ^2^H_non-ex_ = -54.1 ‰). Blanks (empty capsules, crushed) were inserted between each sample to minimize ^2^H carryover and purge the reactor between samples. It should be noted that no chitin reference materials with known δ^2^H_non-ex_ values were available commercially at the time of analysis. Keratin reference materials were used because the fraction of exchangeable hydrogen atoms in keratin (i.e. ∼0.15-0.20) is similar to that of chitin (17, 18). For natural abundance samples with values encompassed by the reference materials, analytical errors were ±2.0 ‰. Highly enriched samples with values outside the range of linear calibration encompassed by the reference materials may have higher errors estimated at ±5-10 ‰. Two water reference materials, USGS47 (δ^2^H= -150.2 ‰) and BARREL-2 (δ^2^H= +800 ‰), were included in every run to monitor slope and isotopic drift.

All ^2^H data is reported in mass-fraction notation, in units of parts per-million (ppm), to accommodate for the gravimetric mass-fraction enrichment method used in the study. Conversion from δ^2^H‰ values (relative to Vienna Standard Mean Ocean Water (VSMOW)) to ppm followed the formula:

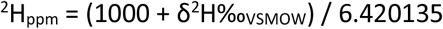

### Statistical analysis

To identify marked (^2^H enriched) specimens, we relied on comparing mosquitoes collected as emerging adults from the enriched larval sites during enrichment with those collected indoors just before enrichment started. During enrichment (October-November), a large gap in ^2^H values (>20 ppm) between the highest value of the pre-enrichment distribution (142 ppm) and lowest value of the marked mosquitoes from enriched larval sites allowed us to classify enriched and non-enriched mosquitoes readily based on ^2^H values (Fig. 1). The first estimator of the fraction of marked mosquitoes was, therefore, the fraction of mosquitoes whose values exceeded 142 ppm. In the subsequent three months (December-February) this gap between the unmarked and the putatively marked mosquitoes decreased as previously documented (11) and (12, 13). Because of the expected “decay” in ^2^H over 4-6 months since marking (11) we hypothesized that certain marked mosquitoes which their ^2^H values decayed enough may accumulate in the upper tail of the population, forming an excess of mosquitoes in that part of the 2H distribution. Accordingly, the proportion of mosquitoes in the post-enrichment population whose ^2^H values were above the 3^rd^ quartile (i.e., the 75^th^ quantile) of the pre-enrichment distribution would be larger than the expected 25%. Our second estimator of the fraction of marked mosquitoes was, therefore, the excess mosquitoes over and above the expected 25%. A conservative statistical test for that hypothesis is testing if the proportion of mosquitoes in the post-enrichment distribution also exceeded the 95% upper confidence limit (UCL) of the 3^rd^ quartile of the pre-enrichment distribution by more than 25% of the ^2^H values. The 95% UCL estimate was derived using the distribution-free method implemented by Proc Univariate (19). For example, if the fraction above the 3^rd^ quartile of the pre-enrichment population was 47%, we attributed the excess 22% (over the expected 25%), to marked mosquitoes, some of which have decayed below the 142 ppm cutoff.

To test the difference between quantiles of pre- and post-enrichment distributions, we used quantile regression implemented by Proc Quantreg (19), which extends the general linear model for estimating conditional change in the response variable across its distribution as expressed by quantiles, rather than its mean (though the median is similar to the mean in symmetric distributions). Quantile regression does not assume parametric distribution (e.g., normal) of the random error part of the model, thus it is considered semi-parametric. The benefit of this analysis is that it addresses low level of enrichment that could be detected in the higher quantiles even when the mean or the median are less affected. The parameter estimates in linear quantile regression models are interpreted as in typical GLM, as rates of change adjusted for the effects of the other variables in the model for a specified quantile (20).

The third method to estimate the fraction of marked mosquitoes in a population, which is independent of a reference subpopulation, was using finite mixed distribution models (FMM). We hypothesized that the post-enrichment population consisted of a subpopulation of natural, non-enriched mosquitoes, and of enriched mosquitoes that had blended into each other following ^2^H loss in the latter (12, 13). Finite mixture models estimate the parameters of the component distributions in addition to the mixing probabilities. Finite mixture models are especially useful for estimating multi-modal or heavy-tailed densities and for classifying observations based on predicted component probabilities (21). These models are increasingly utilized in ecological studies (e.g. mark-capture and migration) (22). We assumed that both subpopulations exhibit a normal distribution of ^2^H values with distinct means and possibly different variances. Analysis was done using PROC FMM, implemented in SAS 9.4 to estimate these parameters of each population and whether there is overall significant support for the blending of these two populations, in which case we used the proportion of the subpopulation with the higher mean as an estimate of marked mosquitoes.

We relied on the three different methods to estimate the fraction of marked mosquitoes, especially after three months or longer post-enrichment when marking decay could have been more influential. Each method has its strengths and assumptions thus the agreement between their results is a measure of the confidence in the overall estimate, which was calculated as the average of all estimators (zeros and negative values included).

We used the inter-quartile range (IQR), calculated as the difference between the 3^rd^ quartile (also known as the 75% quantile) and the 1^st^ quartile (also known as the 25^th^ quantile) as a measure of population spread. The nonparametric skew, defined as (mean – median) / standard deviation was used as a measure of asymmetry (skewness) which is a third of the Pearson 2 coefficient of skewness because it bounds between −1 and +1 for any distribution.

Except in comparisons with distant sites (i.e., Niono and Ballabougou), analysis included desiccated specimens preserved in silica gel. However, some of the specimens from distant sites, were collected for different projects and preserved in 75% ethanol. Preservation in ethanol was found to increase the mean ^2^H value of the mosquitoes by 4.67 ppm (Table S1). Visual comparisons of these distributions were carried out after adjusting for the preservation method (by subtracting 4.67 ppm) for all specimens preserved in ethanol (Fig. 4b and 4d).

## Notes

### Competing Interest Statement

The authors have declared no competing interest.

