## Supplementary material for "Evaluating the contribution of aestivation to the persistence of malaria mosquitoes through the Sahelian dry season using stable isotopes": Fig. S*

**Supplementary Information**

Methods

Although the MRR experiment was carried out in Thierola and M’Piabougou, our focal villages, we included samples from nearby villages: Zanga, Bako, and Dodougou all within 7 km from our focal villages. Additionally, to better understand regional variation in natural ^2^H levels we included samples from Sokourani, located in the Niono rice irrigated area (approximately 60 km long and 5 km wide) ~140 km from the focal villages, as well as Ballabougou, a village located ~25 km north of the focal villages (Fig. S1). Niono is known for its exceptionally high density of *A. colouzzii*. Both this density and its position with respect to the north-easterly wind the prevails during the dry season (December-April) flag Niono as a possible source of windborne long-range migrant mosquitoes that could arrive to Thierola and even beyond (Huestis et al. 2019, Florio et al. 2020). While Niono is the most plausible source, the inner delta of the Niger River near Mopti, which is an additional 150 km northeast may also meet these pre-requisites as possible source of long-distance windborne immigrants. However, no samples have been obtained from the Mopti region.

**Figure S1.** Map of study area including its location (orange rectangle) on a map of Mali (inset). Location of the focal villages (red) surrounded by nearby villages (green) and distant villages (blue) is shown.

Maps we plotted using SAS 9.4 which is licensed to include maps from Gfk GeoMarketing GmbH (inset) and Open Street map.

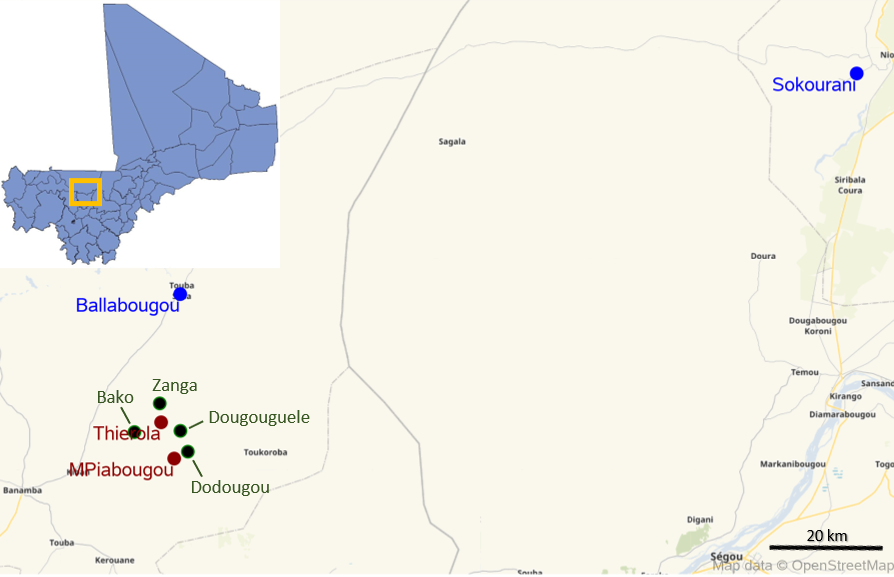

Enrichment larval sites in Thierola and M’Piabougou in relation to the village’s houses are shown in the Fig. S2. In Thierola all larval sites, were sections of a single large pond that flanks the villages from the northwest (Fig. S2, top), except TLV1, which was a separate puddle, and TLV4, TLV7, and TLV10, which were sections from a second large puddle located ~1.2 km northeast of the village.

**Figure S2.** Thierola (top) and M’Piabougou (bottom) aerial images; houses and larval sites.

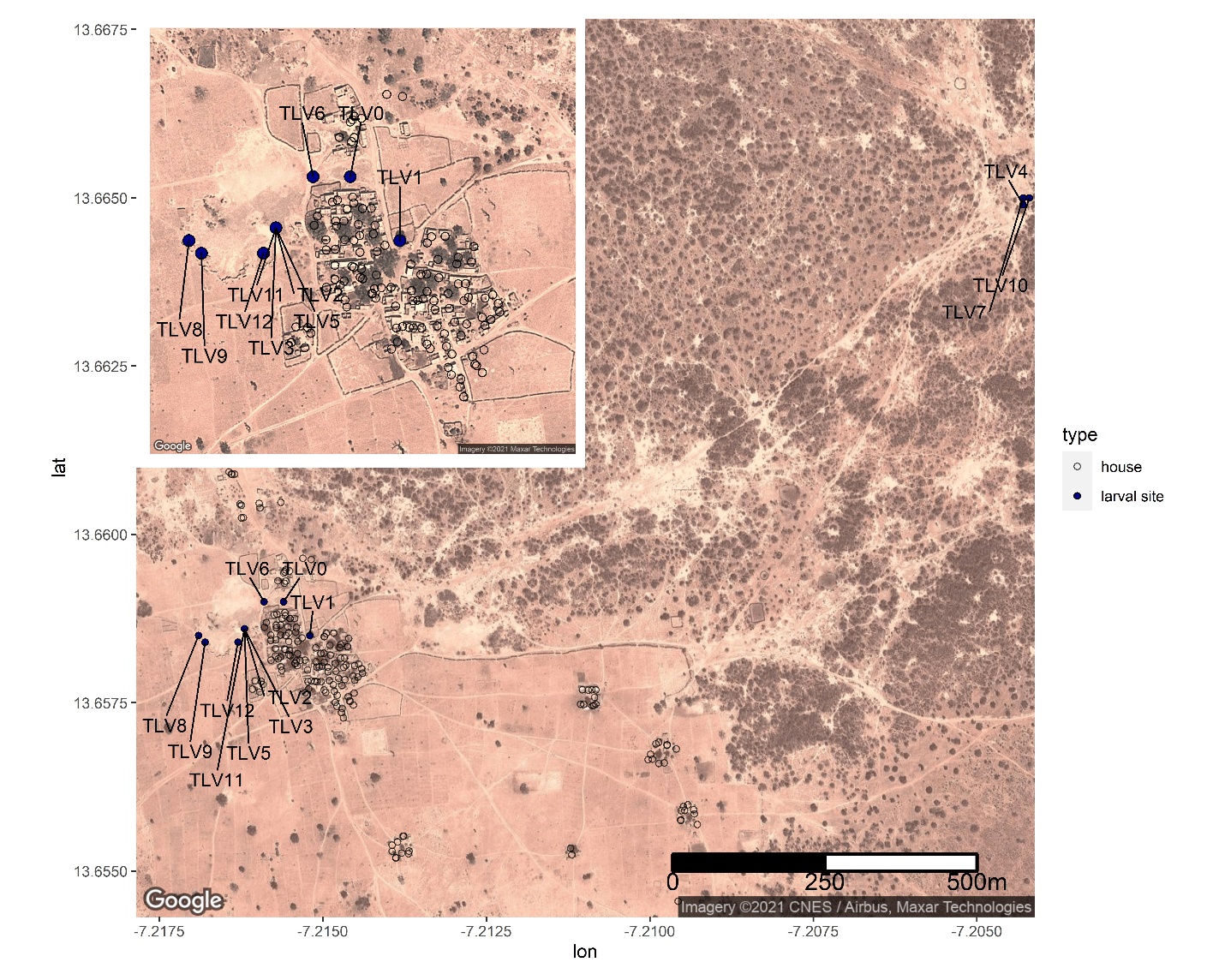

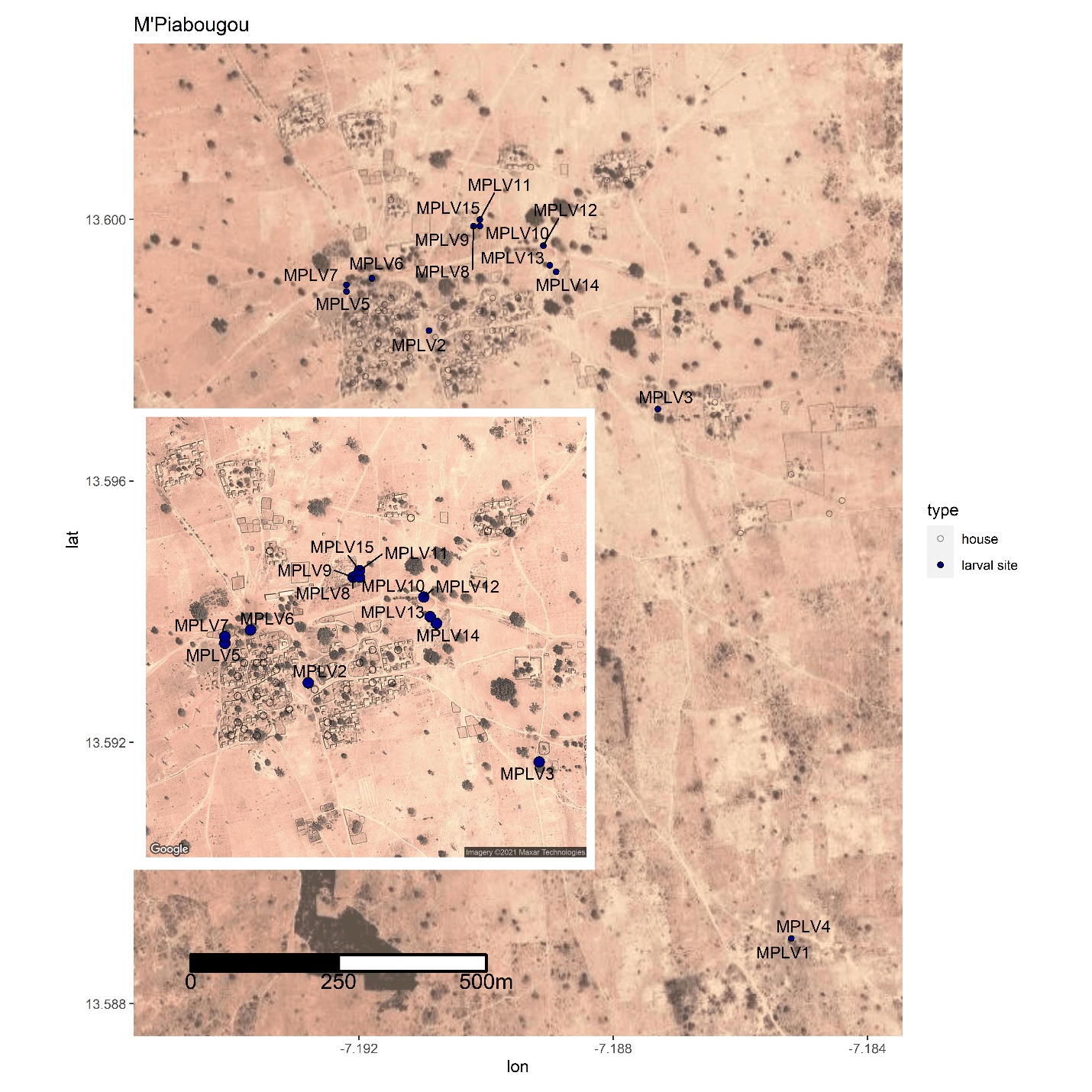

To test if larval sites enrichment had a long-lasting effect beyond the year of treatment, we sampled the first cohort of fourth-instar larvae and pupae from the same larval sites 11 days after the first rains (July 5-21, 2018) and compared their ^2^H levels with those of the previous year (collected indoors in September 2017) as well as fourth-instar larvae and pupae collected during the enrichment experiment from these larval sites (November 2017). As expected, the results revealed no statistical differences in ^2^H levels between pre-enrichment adults and those collected directly from larval sites the following rainy season but large difference between them and the freshly enriched mosquitoes (Fig. S3 and Fig. 1). The results support that the due to evaporation and possibly infiltration the lingering deuterium concentration in the enriched larval sites are negligible.

**Figure S3.** Differences between pre-enrichment mosquitoes from Thierola collected indoors (September 2017: PreEnrContInd), experimentally marked mosquitoes collected as pupae and stage-4 instars from enriched larval sites during enrichment (LV1Enrich and LV5Enrich) and naturally occurring pupae and stage-4 instars collected after the first rain the following year (July 2018: LV1PostEnrich and LV5PostEnrich). Sample size is shown in blue, and green reference lines show the range of natural ^2^H levels as in Fig. 1.

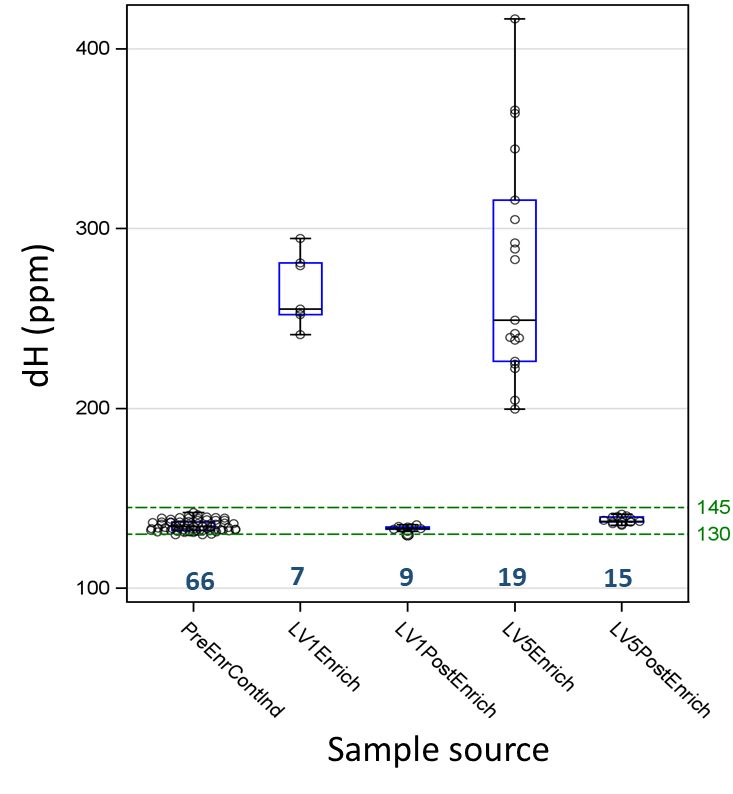

**Figure S4.** Natural temporal variation within and between population in median ^2^H (Y axis), spread of ^2^H as measured by the inter-quartile range (bubble size), and skewness measured by the non-parametric skewness (values inside bubbles; blue denotes negative skewness and yellow and red denote weaker and stronger positive skewness, respectively as seen in scale bar). Sampling years (August to July) are shown on each line and the letters T and M denote samples for Thierola and M’Piabougou.

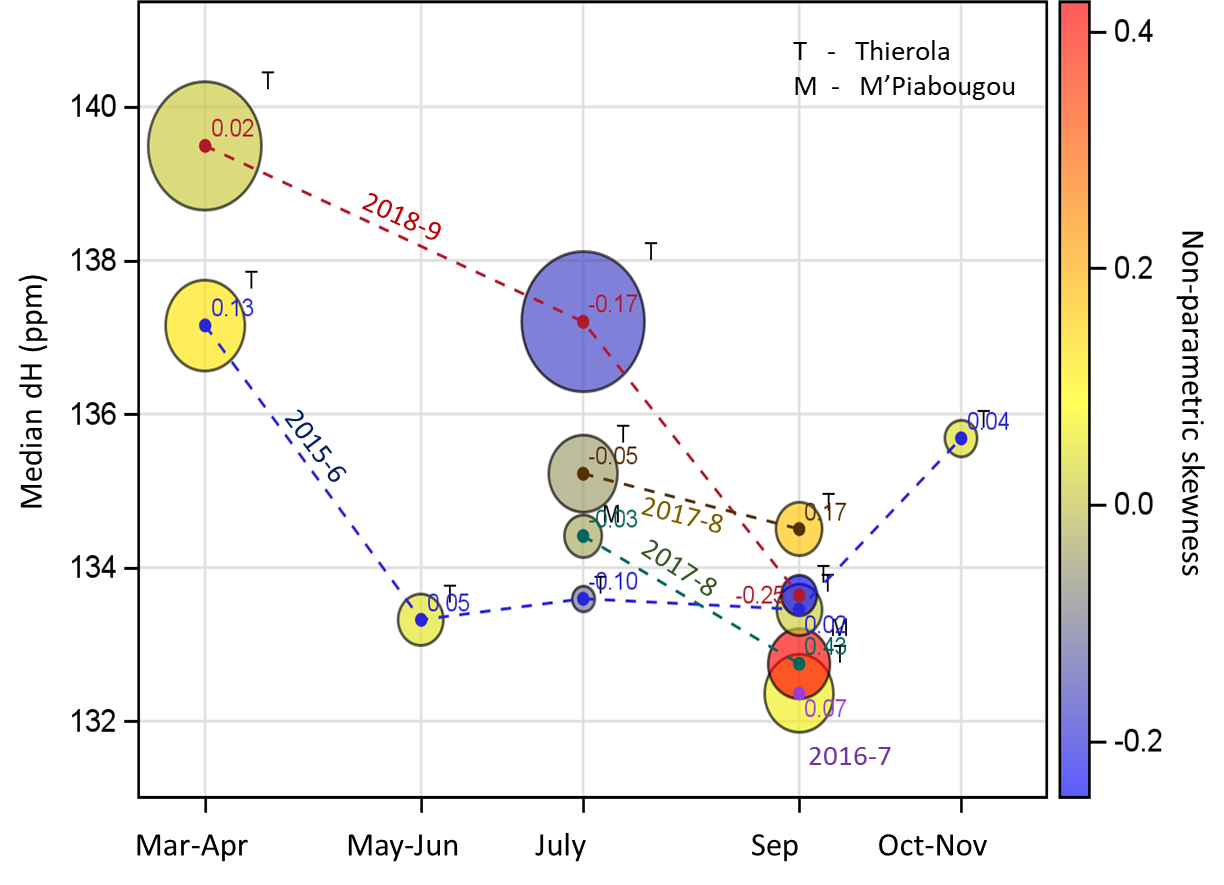

| **Period** | **Population**^1^ | **Enriched (%)^2^** | **3rd quartile**  **Excess (%)**^3^ | **FMM: Enriched**  **Sub-Population (P value)** | **Est. *A. coluzzii* mean** | **Est. Aestivators** |
| --- | --- | --- | --- | --- | --- | --- |
| Late WS (2017) | Thierola | 56.4% (>160 ppm) | ND | ND |  |  |
| Late WS (2017) | M'Piabougou | 10% (>160 ppm) | ND | ND |  |  |
| Late WS (2017) | Pooled | 33% |  |  | 33% | 100% |
| Early DS (2017-8) | Thierola | 33% (>152 ppm) | ND | ND |  |  |
| Early DS (2017-8) | M'Piabougou | 0% (1 mosquito collected) | ND | ND | 29% | 87% |
| Early DS (2017-8) | Pooled | 29% |  |  |  |  |
| Late DS (2018) | Thierola | 24% (>142 ppm), 6.5% (Late DS 2016) | 33%***** (pre-enrich), -2%^ns^ (Late DS 2016) | 19% (F,***)(M, ns) |  |  |
| Late DS (2018) | M'Piabougou | No samples available | No samples available |  |  |  |
| Late DS (2018) | Pooled |  |  |  | 16.5 | 50% |
| Rain Onset (2018) | Pooled | 6.5% (>142 ppm), 0% (Neighbor. Village: 2018) | 21%* (pre-enrich), 8.5%ns (Neighbor. Village: 2018) | 2.5% (***) | 6.1 | 18% |
| ^1^Pooled populations: values reflect means over Thierola and M'Piabougou  ^2^Based on the maximum ^2^H value in the pre-enrichment sample, 142 ppm (see text). No additional tests were needed if ^2^H values of suspected enriched were much larger than the threshold, e.g., 160 ppm. | | | | |  |  |
| ^3^Based on the 3rd quartile of the pre-enrichment or another reference population. Statistical significance is based on excess above the upper 95% CL of the 3rd quartile. | | | |  |  |  |
| WS - wet season, DS - dry season, Pct - percent, ND - not done, F – females, M – males, ns - non-significant, * - P<0.05, *** - P<0.001 | | |  |  |  |  |

**Table S1.** Summary of the estimates of the fraction of mosquitoes with elevated ^2^H over the experiment.
